# NimA promotes cell adhesion at the blood brain barrier of *Drosophila* nervous system

**DOI:** 10.1101/2025.01.15.633121

**Authors:** Rosy Sakr, Sara Monticelli, Claude Delaporte, Gege Zhang, Tarek Tabiat, Angela Giangrande, Pierre B. Cattenoz

## Abstract

Glial cells are crucial for nervous system development and function, by clearing debris, protecting neurons, and ensuring neuronal survival. In Drosophila, glia also form the blood-brain barrier (BBB), which regulates neural stem cell proliferation and shields the nervous system while maintaining organism-wide communication. To uncover glial-specific roles, we compared their transcriptome with that of neurons and macrophages.

Our study identifies NimA, an uncharacterized member of the Nimrod scavenger receptor family, as a glial-specific protein consistently expressed during development. Unlike other family members, i.e. NimC1 (macrophage-specific), Draper (shared by glia and macrophages), and NimC4/Simu (restricted to embryonic stages), NimA is not involved in phagocytosis. Instead, it regulates cell-cell interactions and adhesion, crucial for maintaining the tight septate junctions of the larval BBB. Loss of NimA compromises BBB integrity, delays development, reduces brain size and impairs neural stem cell proliferation.

Altogether, the identification of a novel molecular player and more in general, of the glial-specific molecular landscape, are key to understand the contribution of glia to the construction and the function of the nervous system.

## Introduction

*Drosophila* glial cells play numerous essential roles in the development and in the function of neurons, depending on their position within the nervous system. Typically, perineural (PG) and subperineural glia (SPG) of the central nervous system (CNS) form a tight cellular barrier, fulfilling comparable functions to the vertebrate blood brain barrier (BBB) (Dunton, Gopel et al., 2021). Within the CNS, the cell bodies of neurons and neural-stem cell-like progenitors are insulated and nurtured by the cortex glia (CG) (Hartenstein, 2011). Neurons project dendrites and axons in neuropils that are isolated by the ensheathing glia (EG), while axons in the periphery are ensheathed by wrapping glia (WG) (Hartenstein, 2011, Yildirim et al., 2019). Within the neuropils, the astrocyte-like glia (ALG) send projections to modulate synaptic activities (Stork, Sheehan et al., 2014). At last, CG, ALG and EG have phagocytic functions at various developmental stages (Hilu-Dadia & Kurant, 2020).

Given the multiple roles, including in immunity, it is important to define a glial-specific signature and the molecular features of specific glial subpopulations. Bulk high throughput sequencing technologies raise the resolution of cell definition to genomic scale and provide a global picture of the glial-specific transcriptional landscape. Furthermore, the high sequencing depth and average measurement of gene expression across the glial cell population, makes it possible to identify key actors regulating the multiple roles of distinct glial subpopulations. In this study, we determined the molecular signature of *Drosophila* glia upon comparing their transcriptome to that of cells sharing the same ontogeny but have distinct function, *i.e.* neurons, or have distinct ontogeny but share similar immune and phagocytic functions, *i.e.* macrophages (also called hemocytes). To extrapolate the most robust and stable features, the comparison was carried out at two stages at which the three cells types are fully differentiated but embedded in different environments, late embryonic stage (stage 16, E16) and wandering 3^rd^ larval instar (L3). Embryos represent a closed system in which organogenesis takes place, whereas larvae are constantly exposed to metabolic as well as immune challenges and at that stage takes place the growth and development of the adult tissues. These data allow us to describe robust glial-specific features that are stable during development. We identify Nimrod A (NimA), which belongs to the family of phagocytic receptors Nimrod (Kurucz, Markus et al., 2007, Somogyi, Sipos et al., 2008). Unlike the other members of the Nimrod family, NimA does not seem to be involved in the regulation of phagocytosis. Instead, we demonstrate that NimA mediates cell aggregation and that its loss results in defective BBB as well as altered neural development.

Is sum, we identify the glial-specific transcriptional landscape and unveil NimA as a novel transmembrane protein required to build an efficient cellular barrier.

## Results

### Transcriptomic signature of glia

To determine the molecular signature of glia, we identified the genes upregulated in these cells compared to neurons and macrophages at both E16 and L3, allowing us to define a robust glial-specific signature **(Figure 1A, Supplementary Figure S1A,B** for genes upregulated in neurons and macrophages, respectively). This comparison highlights known markers that reflect glia origin and function, including the glial cell fate determinant Glial cell missing (Gcm) (Hosoya, Takizawa et al., 1995, Vincent, Vonesch et al., 1996), the panglial transcription factor Reversed polarity (Repo)(Halter, Urban et al., 1995), the ALG marker Nazgul (Naz) (Ryglewski, Duch et al., 2017), the G-protein coupled receptor Moody regulating BBB functions (Bainton, Tsai et al., 2005), the WG marker Nervana 2 (Nrv2) (Stork et al., 2008). This validates our approach and also allows us to define new glial markers such as the predicted transmembrane proteins (CG1537, CG1545) and a sugar transporter (CG8837) **(Supplementary Figure S1C)**.

**Figure 1:**
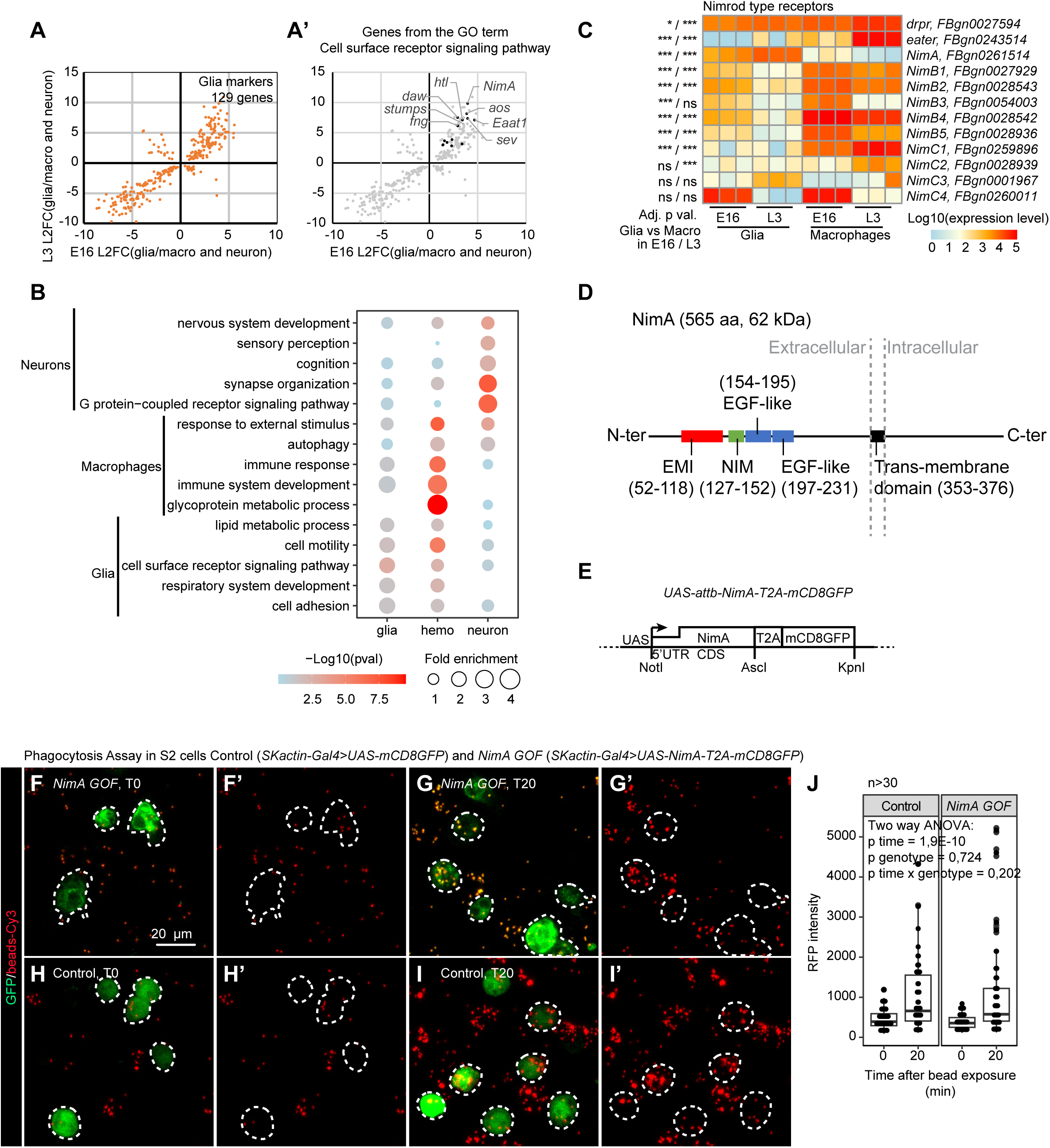
Identification of the molecular signature of glia. **A,A’**) Comparison of the glia transcriptome to the neurons’ and macrophages’ transcriptomes from stage 16 embryos (E16) and wandering 3^rd^ instar larvae (L3). The scatter plot represents the Log2 fold change (L2FC) of differentially expressed genes enriched in glia compared to neurons and macrophages. Only genes presenting significant fold change in both E16 and L3 were plotted, the x-axes represent the L2FC in E16 and the y-axes the L2FC in L3. The genes annotated with the GO term “Cell surface receptor signaling pathway” are highlighted in black in (**A’**). The data are available in **Supplementary Table S1**. **B)** GO term enrichment analyses on the markers specific to neurons, to macrophages and to glia. The 5 most significant GO terms are represented here. The size of the dots represents the enrichment levels of the GO term and the color gradient, the p-value (-Log10(pval), from light blue to red for low to high significance). **C)** Heatmap representing the expression levels of the Nimrod transmembrane receptors in macrophages and glia. The color gradient is representative of the expression levels from blue to red, low to high expression. Note that most Nimrod receptors are enriched in macrophages, only few are enriched in glia (*i.e.* NimA, NimC4). The adjusted p-values mentioned on the left column were estimated using Wald test (using DESeq2) and are issued from the comparison of the expression levels in glia and macrophages at E16 or L3 (p E16 / p L3), values in **Supplementary Table S1.** **D)** Structure of the glial-specific receptor NimA. **E)** Schematic representation of the expression vector *UAS-attb-NimA-T2A-mCD8GFP*. The vector expresses NimA under the UAS promoter. The self-cleaving peptide T2A followed by the membrane tethered GFP (T2A-mCD8GFP) is in frame with NimA. Thus, the translation of *NimA-T2A-mCD8GFP* mRNA produces two proteins, *i.e.* NimA and mCD8GFP to allow the tracking of NimA expressing cells with the GFP. to track transfected cells with the expression of mCD8GFP. **F-J)** Phagocytosis assay on S2 cells exposed to fluorescent latex beads. S2 cells transfected with *NimA* expression vector (*UAS-attb-NimA-T2A-mCD8GFP*) or control vector (*UAS-mCD8GFP*) were challenged with latex beads (in red) for 0 min (T0) or 20 min (T20) and analyzed by confocal microscopy. Full stacks are displayed, scale bar is 20 µm, the transfected cells are surrounded by a white dashed line, (**F-G)** are NimA overexpressing cells (gain of function or NimA GOF) at T0 (**F**) and T20 (**G**) and (**H-I**) are controls, red channel only are shown in (**F’,G’,H’,I’**). (**J**) Boxplot quantifying the RFP intensity at T0 and T20 in control and NimA GOF S2 cells. N>30, p-values were measured by two-way ANOVA with the variables time and genotype.

Gene Ontology (GO) analyses of the cell-specific signature returned expected terms such as immune response for macrophages and synapse organization for neurons, reflecting the well-known role of these cell types in the defense from the non-self and in cell circuitry, respectively (**Figure 1B**). Interestingly, the GO analyses reveal a different behavior for glia, as we found lower enrichment for these cells. This reflects the fact that specialized glial subtypes perform distinct functions: phagocytosis for ALG, CG and EG, insulation for EG, WG and SPG (**Figure 1B, Supplementary Table S2**). The most significant GO term enriched in glia is cell surface receptors, in line with glial cells being in constant interaction with neurons (**Figure 1B**). Amongst the first ten genes identified by this GO (**Supplementary Table S2**), the Nimrod A (NimA) transmembrane protein (**Figure 1A’,D**) belonging to the family of Nimrod scavenger receptors, displays the strongest enrichment in glia.

In sum, our analysis identifies stable cell-specific transcriptomic features that define glia throughout development and highlights NimA as a novel glial-specific marker.

### The glial-specific cell surface protein NimA does not act as a scavenger receptor

In *Drosophila*, the Nimrod family contains 12 members defined by the presence of specific NIM and EGF-like repeats (Kurucz et al., 2007, Somogyi et al., 2008). Most Nimrod proteins that have been characterized are involved in cell debris or pathogen recognition to promote phagocytosis. They include the well-studied phagocytic receptors Eater and NimC1 involved in the phagocytosis of pathogens by macrophages (Pearson et al., 1995; Rämet et al., 2001) as well as Draper (Drpr) and Nimrod C4 (NimC4) that act in macrophages and glia to promote the clearance of apoptotic cells and axonal debris (Awasaki, Tatsumi et al., 2006, Kurant, Axelrod et al., 2008, Manaka, Kuraishi et al., 2004). NimA is the only Nimrod protein specific to glia during development (**Figure 1C, Supplementary Figure 1D**).

NimA is a Drpr-like protein containing one NIM domain followed by two EGF-like repeats and one Emilin (EMI) domain (**Figure 1D**) (Callebaut, Mignotte et al., 2003, Kurucz et al., 2007). Its closest mammalian orthologs are Platelet Endothelial Aggregation Receptor 1 (PEAR1) and Multiple EGF-like domains 10 (MEGF10) that contain multiple EGF domains and one EMI domain (DIOPT, (Hu, Flockhart et al., 2011)). Both PEAR1 and MEGF10 regulate glia mediated phagocytosis (Iram, Ramirez-Ortiz et al., 2016, Wu, Bellmunt et al., 2009), hinting to a functional conservation in phagocytosis between NimA and the two orthologs.

To test for the role of NimA in phagocytosis, we monitored the ability of S2 cells to phagocytose latex beads. This straightforward approach has been used to validate the phagocytic properties of NimC1, which promotes binding of macrophages to latex beads (Melcarne, Ramond et al., 2019b). Since NimA is not expressed endogenously in S2 cells (Brown, Boley et al., 2014), we generated a NimA expression vector under the UAS promoter and inserted a T2A-GFP construct in frame with the NimA coding sequences to track the transfected cells with the GFP (**Figure 1E**)(Gonzalez, Martin-Ruiz et al., 2011). Following transfection, cells were challenged with fluorescent latex-beads and bead uptake was estimated by measuring fluorescence intensity at 0 and 20 min after challenge. No significant difference was observed between control and NimA overexpressing cells (**Figure 1F-J**), suggesting that NimA is not regulating phagocytosis or at least, not through the same mechanism as NimC1.

Next, we evaluated the role of NimA in glia phagocytosis *in vivo*. A classic example of neuronal remodeling mediated by glia is the phagocytosis of the Pigment-dispersing factor-tri neurons (PDF-tri neurons) that appear at mid-pupal stage and undergo apoptosis within three days following hatching (**Figure 2C**) (Helfrich-Forster, 1997, Vita, Meier et al., 2021). PDF-tri neuron removal requires the scavenger receptor Drpr in the EG and CG wrapping the area (Vita et al., 2021). Available single cell RNA sequencing (scRNAseq) data on the adult brain show that NimA is detected in EG and CG (**Figure 2A,A’**) (Davie, Janssens et al., 2018), suggesting that it may act along Drpr to promote PDF-tri neuron phagocytosis.

**Figure 2:**
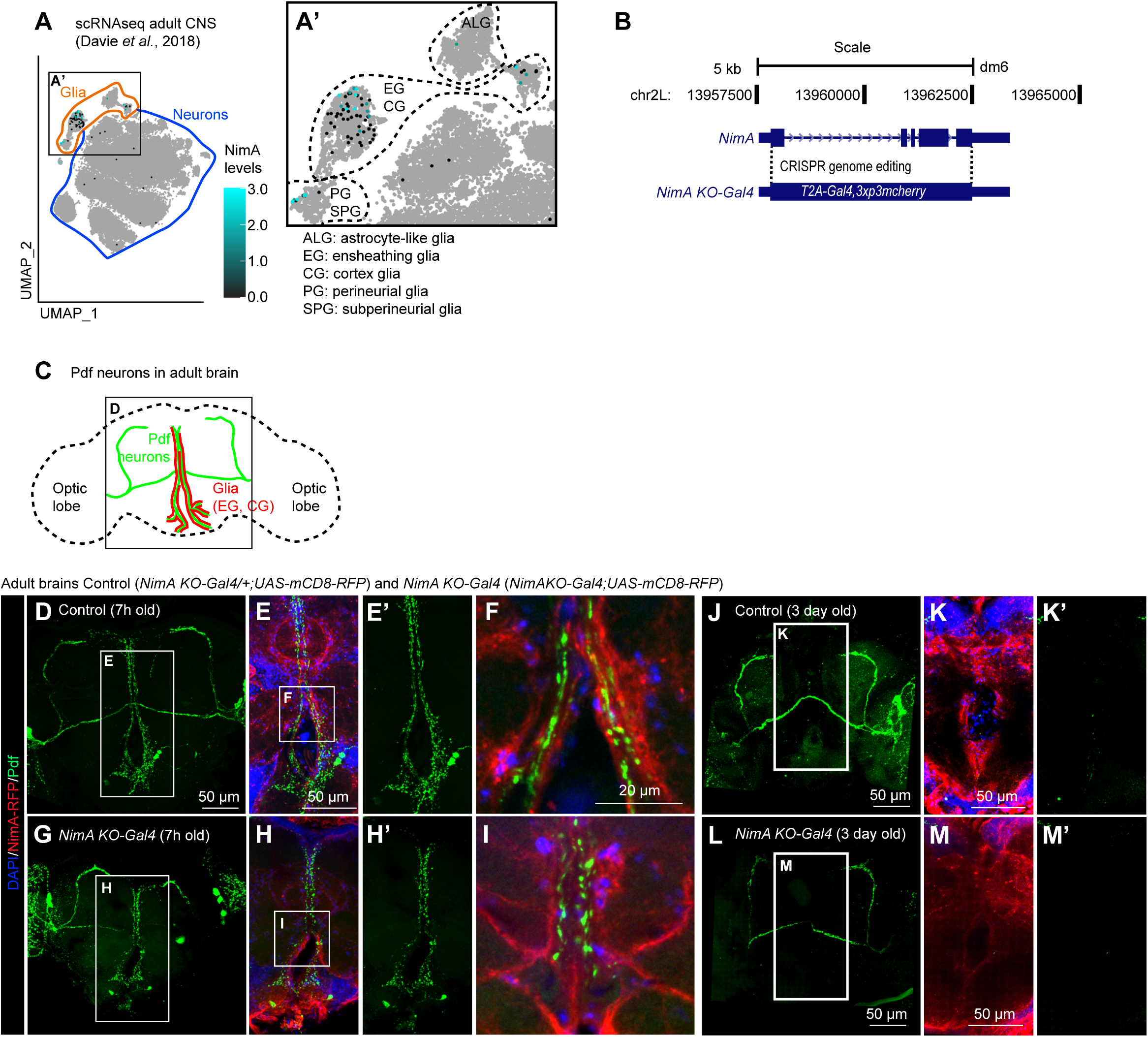
The scavenger receptor NimA is expressed specifically in glia. **A,A’)** UMAP representing *NimA* expression profile in scRNA seq data from adult CNS (data and annotation from (Davie et al., 2018)). *NimA* expression levels are represented with color gradient from black to blue. Higher magnification around the glia cluster is shown in (**A’**) with annotation for the perineural (PG) and subperineural glia (SPG), the astrocyte-like glia (ALG), the ensheathing glia (EG) and the cortex glia (CG). **B)** Schematic representation of the *NimA-T2A-Gal4* allele (*NimA KO-Gal4*). The coding sequence of *NimA* was replaced by *T2A-Gal4*, producing a null allele that serves also as a *Gal4* reporter. **C)** Schematic representation of the PDF-tri neurons surrounded by glia in the adult CNS. **D-M)** Immunolabelling of *NimA* heterozygous (*NimA-T2A-Gal4/+;UAS-mCD8RFP*) or *NimA KO-Gal4* homozygous (*NimA-T2A-Gal4;UAS-mCD8RFP)* adult brains, 7 h old (**D-I**) or 3-day-old (**J-M**) with anti RFP (in red) and anti-Pdf (in green) that labels the PDF-tri neurons undergoing apoptosis within 3 days after eclosion. (**D,G,J,L**) show full stacks of the brains, (**E,H,K,M**) are substacks at higher magnification of the region covering the PDF-tri neurons, the Pdf channels alone are shown in (**E’,H’,K’,M’**). Single sections at higher magnifications are shown in (**F**,**I**) highlighting the close proximity of *NimA* glia around the PDF-tri neurons. Each immunolabeling was replicated (n>5). Scale bars are 50 µm and 20 µm, as indicated.

To validate NimA expression in the glia surrounding the PDF-tri neurons, we generated a *NimA* knock out allele that also acts as a driver (*NimA KO-Gal4*) upon replacing the *NimA* coding sequence by *T2A-Gal4* using *CRISPR/Cas9* mediated gene replacement (**Figure 2B, Supplementary File S1**). By preserving fully the genomic regulation of *NimA* expression, this strategy produced a faithful *NimA Gal4* driver. *NimA KO-Gal4* crossed with UAS-fluorescent reporters revealed NimA expression in glia surrounding neuropils (**Supplementary Figure S2A-C’’’**) as well as around the PDF-tri neurons in brains from young adults (**Figure 2D-I’**). In 3-day-old adults, the PDF-tri neurons have been cleared in both Control and *NimA KO-Gal4* homozygous animals (**Figure 2J-M’**), indicating that NimA has no impact on the phagocytosis of the PDF-tri neuron debris.

Overall, these data show that NimA is expressed in glia forming boundaries (EG and CG) and suggest that NimA does not function as a scavenger receptor like other members of the Nimrod family.

### NimA is expressed in surface glia of the larval brain and is required for BBB integrity

In addition to their roles in phagocytosis, the scavenger receptors Eater and NimC1 promote cell adhesion and spreading in macrophages (Melcarne et al., 2019b). Provided the expression profile of NimA in ensheathing glia, we hypothesized that similarly to NimC1 and Eater, NimA plays a role in the adhesion between glial cells. This phenotype being more easily observed at an earlier stage than in the adult, we shifted to the larval CNS to test this hypothesis.

The ventral nerve cord (VNC) of the larval CNS contains two longitudinal neuropils fully wrapped by EG (**Figure 3A,C**). *NimA KO-Gal4* shows expression in Repo positive glia in the CNS but not in the peripheral nervous system (PNS) (**Figure 3B, Supplementary Figure S3B-F’’’**). The expression profile of *NimA KO-Gal4* is concordant with fluorescent *in situ* hybridization (FISH) data obtained using a *NimA* probe. NimA is expressed in the EG isolating the neuropils as well as in the SPG of the larval CNS (**Figure 3B, Supplementary Figure S3A-F’’’**).

**Figure 3:**
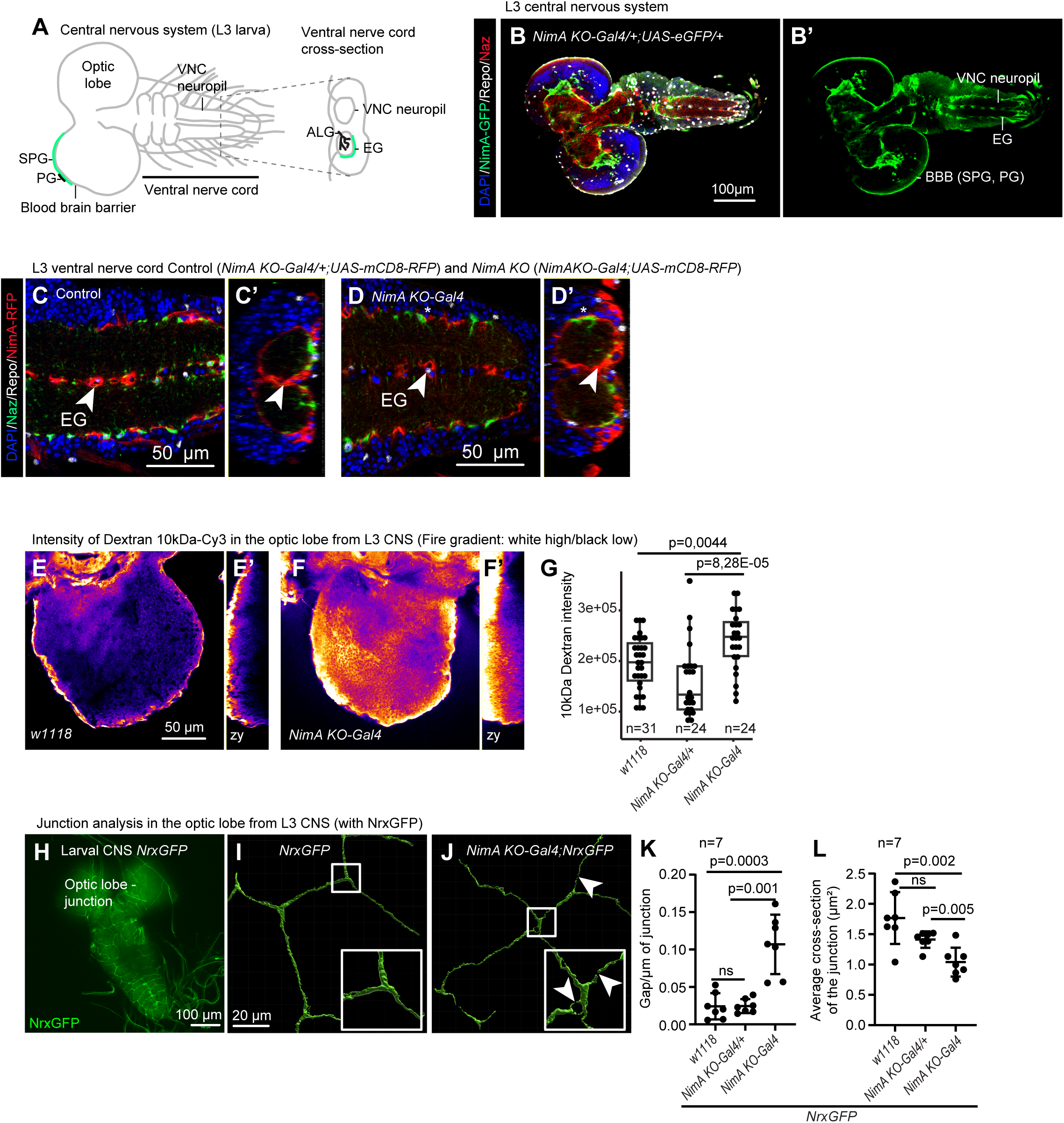
*NimA* is expressed in glia forming barriers. **A)** Schematic representation of the larval CNS. The perineural (PG) and subperineural glia (SPG), the astrocyte-like glia (ALG) and the ensheathing glia (EG) are shown on the schematic. The SPG and EG expressing NimA are indicated in green. **B,B’**) Immunolabelling of larval CNS *NimA-T2A-Gal4/+;UAS-eGFP/+* with anti-Nazgul labelling ALG (in red), anti-Repo labelling all glia (in white) and anti-GFP labeling *NimA* cells (in green). Single sections of the larval CNS with all channels or the GFP channel are shown in (**B’**). DAPI was used to label nuclei (in blue) and the scale bar represents 100 µm. **C,D**) Immunolabelling of larval ventral nerve cords from *NimA-T2A-Gal4/+;UAS-mCD8RFP/+* (Control, **C**) or *NimA-T2A-Gal4;UAS-mCD8RFP/+* (*NimA KO-Gal4*, **D**) animals with anti-Nazgul (in green), anti-Repo (in white) and anti-RFP (in red). Single sections are shown with all channels in (**C,D)**. Cross-sections at levels of the arrowheads highlight the ensheathing glia wrapping the neuropils (**C’,D’**). DAPI was used to label nuclei (in blue) and the scale bar represents 50 µm, each immunolabeling was replicated (n>8). **E-G)** Penetration assay carried out on wild type (*w1118*, **E,E’**), *NimA KO-Gal4/+* (*NimA-T2A-Gal4/+*) and *NimA KO-Gal4*(*NimA-T2A-Gal4*, **F,F’**) larval CNS with 10 kDa fluorescent Dextran. The optic lobes of the CNS were acquired by confocal microscopy, single sections localized in the middle of the optic lobes are shown in (**E,F**), the Fire color gradient is proportional to the fluorescence intensity from black (low intensity) to white (high intensity). Cross-sections are shown in (**E’,F’**). The scale bar represents 50 µm. The intensities in the optic lobes are quantified in (**G**), p-values were estimated by ANOVA (p ANOVA = 7,4E-05) with post-hoc student test. **H-L)** Analysis of the septate junctions with Neurexin-GFP (NrxGFP). The fusion protein NrxGFP marks the septate junctions formed by the subperineural glia (SPG) in the L3 CNS (**H**). (**I,J**) 3D reconstruction of the SPG septate junctions from the optic lobes of *NrxGFP* (**I**) and *NimA KO;NrxGFP* (**J**) larvae. Gaps in the septate junctions are indicated by white arrowheads. The number of gaps normalized by the length of the junctions (Gap per µm) is shown in (**K**) and the thickness of the junction (average cross-section) in (**L**). N=7, the p-values were estimated with student tests.

No gap nor wrapping defects were detected in the EG surrounding the neuropils of the VNC in *NimA KO-Gal4* homozygous larvae (**Figure 3C’,D’**). No defects could be detected either in the EG wrapping the neuropils of the brains (data not shown).

We then monitored the integrity of the larval BBB, which is composed of PG and SPG. The BBB loses integrity when the septate junctions of the SPG are altered (Stork et al., 2008). We estimated BBB permeability by incubating the CNS in 10 kDa dextran coupled with a fluorescent dye that cannot cross the BBB in normal conditions. We found a significant increase in dextran intensity within the CNS of *NimA KO-Gal4* homozygous compared to heterozygous and wild type larvae (**Figure 3E-G**). Then, to assess the integrity of the BBB septate junctions, we used the Neurexin-GFP (NrxGFP) transgenic line expressing a fusion protein between GFP and Neurexin IV, which localises to septate junctions (Baumgartner, Littleton et al., 1996, Laval, Bel et al., 2008) (**Figure 3H-J**). *NimA KO-Gal4* homozygous CNS display thinner septate junctions with multiple gaps compared to heterozygous and wild type CNS (**Figure 3K,L**).

These data show that NimA is required to build and/or maintain integral septate junctions and preserve the impermeability of the BBB.

### NimA is required for normal neural development

The integrity of the BBB is essential for CNS development and its alteration is associated with a dysregulation of neural stem cell proliferation in the larva (Speder & Brand, 2014). The *NimA KO-Gal4* homozygous larvae show reduced CNS size in L3 (**Figure 4A-C**). This phenotype does not seem to rely on increased cell death, as the number of apoptotic cells observed upon immunolabeling with the Dcp1 marker (Song, McCall et al., 1997) is similar to that observed in control animals (**Figure 4D**). By contrast, cell proliferation is decreased in CNS from *NimA KO-Gal4* homozygous larvae, which associates with a reduced pool of neural stem cells (**Figure 4E,F**) and likely accounts for the undersized CNS. Consistently with the observed CNS defects, *NimA KO-Gal4* homozygous larvae display a significant pupariation delay and increased lethality rate by the end of the larval stage (**Figure 4G,H**). No lethality nor further delay were observed at eclosion and the lifespan of *NimA KO-Gal4* homozygous flies is comparable with the one of control animals (**Figure 4I-K**). Finally, mutant animals are fertile and display curved wings in adults, which is comparable to the phenotype of known *NimA* null alleles such as *jaunty^1^*, *NimA^SF7^* and *NimA^j-676^* (Woodruff & Ashburner, 1979).

**Figure 4:**
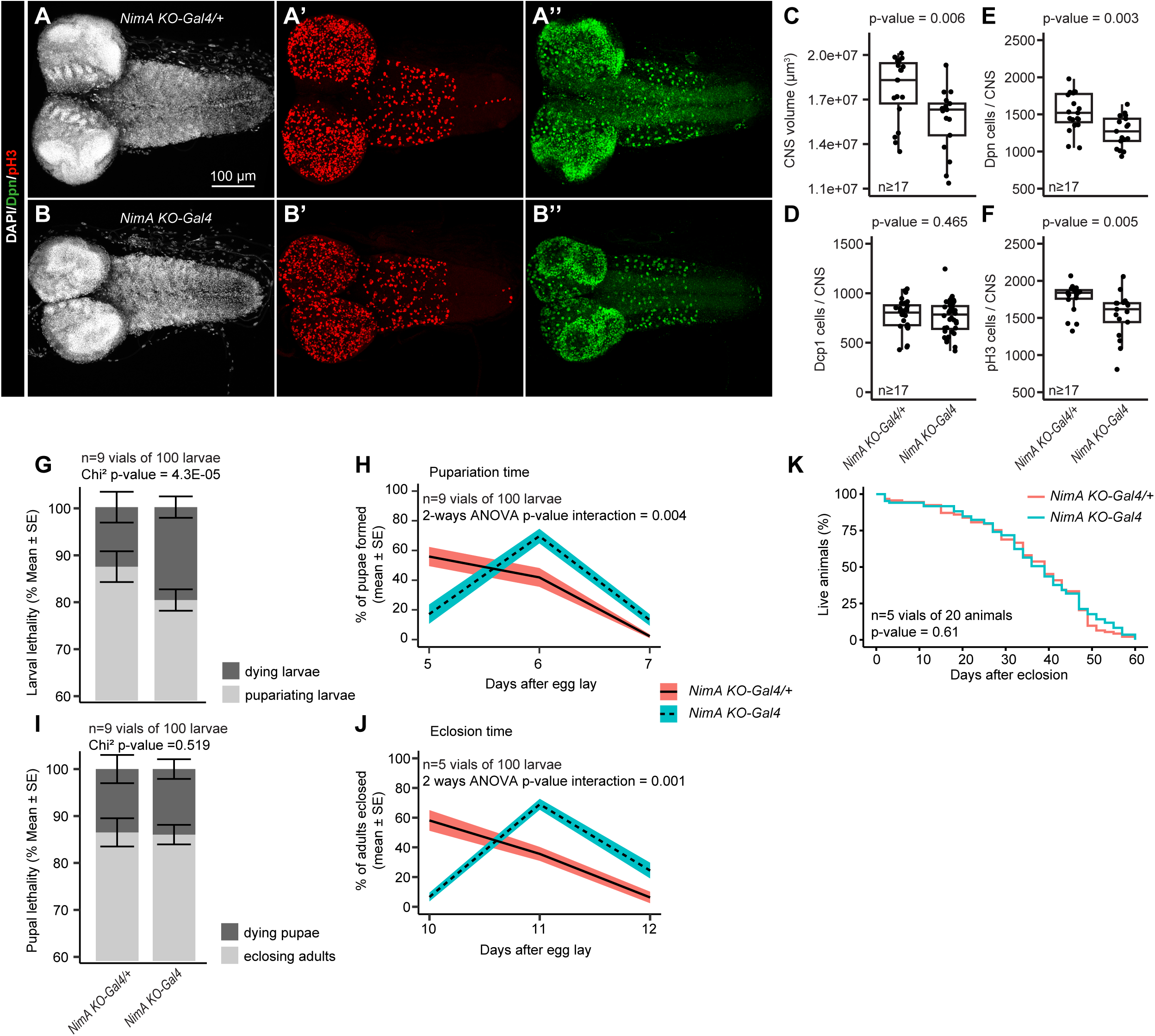
Lack of NimA alters the development of the brain. **A,B)** Immunolabelling of L3 CNS *NimA KO-Gal4/+* (**A**) and *NimA KO-Gal4* (**B**) with antibodies targeting the neural stem cell marker Deadpan (Dpn, in green) and the mitotic marker phospho histone H3 (pH3, in red). Nuclei were labelled with DAPI (in gray). The scale bar represents 100 µm. **C-F)** Quantification of the CNS volume (**C**), the number of apoptotic cells marked with anti-Dcp1 per CNS (**D**), the number of neural stem cell marked with anti-Dpn per CNS (**E**) and the number of mitotic cells marked with anti-pH3 per CNS in *NimA KO-Gal4/+* and *NimA KO-Gal4*L3 larvae. N>=17, p-values were estimated by Mann-Whitney U test. **G-J)** Quantification of lethality at larval stage (**G**), of pupariation time (**H**), of lethality at pupal stage (**I**) and of the imago hatching time (**J**) in *NimA KO-Gal4/+* and *NimA KO-Gal4* animals. P-values were estimated by Chi² test in (**G,I**) and by two-ways ANOVA in (**H,J**). **K**) Quantification of the life-span in *NimA KO-Gal4/+* and *NimA KO-Gal4* animals. The p-value was estimated by log-rank test.

In sum, NimA alteration of BBB integrity is accompanied by a developmental delay of the CNS.

### NimA promotes cell aggregation and forms multimers

Based on the role of NimA in cells forming barriers, we speculated that this protein is involved in cell adhesion. We tested this hypothesis in S2 cells transfected with a *NimA* expression vector containing either a Myc or a HA tag in the C-terminal intracellular domain (**Figure 5A**). NimA accumulates at the cytoplasmic membrane and cytoplasmic projections (**Figure 5B-C**) and the cells expressing NimA tend to aggregate and form large clumps (**Figure 5B-D**).

**Figure 5:**
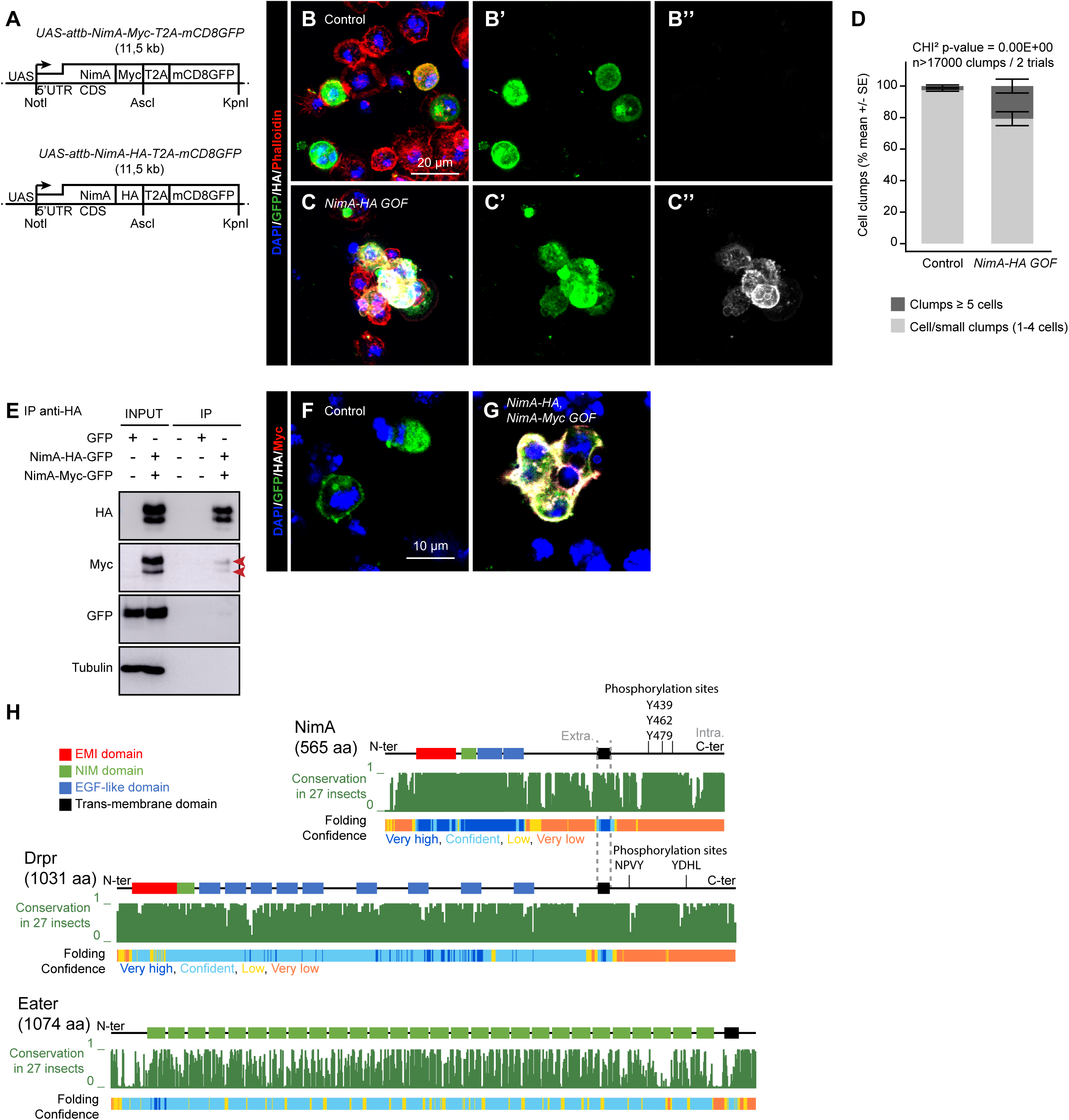
NimA forms homomers and promotes the aggregation of cells. **A)** Schematic representation of the expression vectors *NimA-Myc* (*UAS-attb-NimA-Myc-T2A-mCD8GFP*) and *NimA-HA* (*UAS-attb-NimA-HA-T2A-mCD8GFP*). **B-C)** Immunolabelling of S2 cells transfected with control vector (*UAS-mCD8GFP*, **B-B’’**) or *NimA-HA* expression vector (*UAS-attb-NimA-HA-T2A-mCD8GFP*, **C-C’’**) labelled with anti-GFP (in green) and anti-HA (in white) antibodies and phalloidin (in red). Nuclei were labelled with DAPI. The scale bar represents 20 µm. The images represent full stacks from confocal acquisitions. **D)** Quantification of the number of large clumps in control and *NimA-HA GOF* S2 cells, based on the number of nuclei per clump (>5 nuclei). N>17000 clumps in 2 independent trials, p-value was estimated with Chi² test. **E)** Co-immunoprecipitation of NimA-HA and NimA-Myc in S2 cells transfected by the two vectors described in **(A)**. The immunoprecipitation (IP) was carried out with anti-HA antibody and HA, Myc, GFP and Tubulin were detected in the input and in the IP by western-blot. The two first lanes represent the input for the control (GFP expression vector) and the co-transfected cells (vectors *NimA-HA* and *NimA-Myc*), the three following lanes represent IPs on non-transfected S2 cells, control and co-transfected cells, respectively. **F-G)** Immunolabelling of S2 cells transfected with control vector (*UAS-mCD8GFP*, **F**) or co-transfected with *NimA-HA* and *NimA-Myc* expression vectors (**G**) labelled with anti-GFP (in green), anti-HA (in white) and anti-Myc (in red) antibodies and DAPI (in blue). The scale bar represents 10 µm. The images represent single sections from confocal acquisitions. **H)** Conservation and structure prediction of NimA, its paralog Drpr and the Nimrod receptor Eater. The conservation histograms (in green) across 27 insect species were drawn from the UCSC genome browser track Multiz Alignment & Conservation on the *Drosophila* genome Dm6. The protein domain annotations and folding prediction were taken from Uniprot and alphaFold protein structure database on the proteins NimA (Q8IP58), Drpr (Q9W0A0) and Eater (Q9VB78). Tyrosine phosphorylation sites are indicated in the intracellular domains of NimA and Drpr. For NimA, the sites were predicted with GPS 6.0 (prediction score > 0.99) (Chen, Zhang et al., 2023).

Noteworthily, NimA ortholog PEAR1 is described as a platelet-platelet contact receptor that stabilizes the platelet aggregate upon activation through multimerization (Kauskot, Di Michele et al., 2012, Nanda, Bao et al., 2005). To assess the ability of NimA to form multimers like its ortholog, we carried out an immunoprecipitation assay targeting the HA tag on S2 cells co-transfected with both HA and Myc vectors or T2A-mCD8GFP as a control. The western blot shows that NimA-HA co-precipitates with NimA-Myc (**Figure 5E**). Concordantly, assays on co-transfected S2 cells show colocalization of NimA-HA and NimA-Myc at the cell membrane (**Figure 5F,G**). These data show that NimA is a transmembrane protein that forms multimers and promotes cell-cell adhesion similarly to its ortholog PEAR1.

## Concluding remarks

Our transcriptomic approach identified a panel of 129 genes constitutive of a robust molecular signature for glia preserved during development. Amongst the glial-specific markers, the NimA transmembrane protein is a paralog of the scavenger receptor Drpr. We show that NimA is not involved in phagocytosis, rather, it is expressed in glia forming boundaries such as the SPG and the EG and is required to establish/maintain strong junctions between SPG in the BBB. NimA promotes cell-cell contact and forms homomers like its human ortholog PEAR1. Since PEAR1 multimerization requires the EMI extracellular domain (Kauskot et al., 2012), which is the main domain conserved between both orthologs, we speculate that NimA multimerization is mediated by its EMI domain.

The Nimrod proteins are characterized by single or multiple copies of the NIM and EGF-like domains that define their affinity to specific ligands (Fujita et al., 2012, Kurucz et al., 2007, Melcarne, Lemaitre et al., 2019a, Melcarne et al., 2019b) (**Figure 5H**). Drpr contains up to 15 EGF-like domains (depending on the isoform) and promotes phagocytosis after binding multiple ligands including Pretaporter and phosphatidyl serine exposed by apoptotic cells (Kuraishi, Nakagawa et al., 2009, Tung, Nagaosa et al., 2013) as well as bacteria (Shiratsuchi, Mori et al., 2012). Eater and NimC1, involved in phagocytosis, have extensive extracellular domains with a high number of NIM/EGF-like domains (Kurucz et al., 2007) (**Figure 5H**). The EGF/NIM repeats defines the affinity of the receptors to distinct bacteria. Nimrod genes’ organisation as a cluster is conserved in insects. Specie-specific diversity is observed at the levels of the EGF/NIM repeat numbers and sequences for each gene, suggesting a selection pressure to preserve this cluster adapted for each species (Somogyi, Sipos et al., 2010, Somogyi et al., 2008). Within the Nimrod cluster, NimA displays the shortest extracellular domain with only one NIM and two EGF-like domain (Kurucz et al., 2007) and strong conservation across species (Somogyi et al., 2010, Somogyi et al., 2008) (**Figure 5H**). This suggests that NimA underwent exaptation to shift from an immune function toward a developmental function such as glia-glia adhesion. The high conservation of NimA extracellular domain also suggests specific conserved ligands activating the receptor.

Finally, while the NimA extracellular domain is highly structured, the intracellular domain displays no known protein domain nor conserved feature or structure that could allow inference on its molecular function (**Figure 5H**). Following ligand binding, NimA paralog Drpr is phosphorylated on tyrosine residues in the intracellular domain, which is required for the regulation of phagocytosis (Fujita, Nagaosa et al., 2012, Ziegenfuss, Biswas et al., 2008). Since several conserved tyrosine residues are also predicted to be phosphorylated in the intracellular domain of NimA (**Figure 5H**), it is possible that its multimerization leads to tyrosine phosphorylation to activate the downstream cascade responsible for cell aggregation.

In sum, our transcriptomic study identifies a novel and unexpected role for an uncharacterized member of the Nimrod protein family, helping us understanding the role and biology of glial cells.

## Materials and methods

### Fly strains and genetics

Flies were raised on standard medium at 25°C. Fly strains used are listed in the (**Table 1**). *NimA KO-Gal4* null mutant was generated by CRISPR-mediated mutagenesis and performed by WellGenetics Inc. using modified methods from (Kondo and Ueda, 2013) (**Supplementary file S1**). In brief, the upstream gRNA sequence ACTGCTCCTCCTGCTTGCAA[TGG] and downstream gRNA sequence GTGCCCTTCTAACATATACC[AGG] were cloned into U6 promote plasmid(s). Cassette *Gal4-3xP3-RFP*, which contains ribosome binding sequence (RBS), Gal4, SV40 polyA terminator and a floxed 3xP3-RFP, and two homology arms were cloned into pUC57-Kan as donor template for repair. *NimA*-targeting gRNAs and hs-Cas9 were supplied in DNA plasmids, together with donor plasmid for microinjection into embryos of control strain *w[1118]*. F1 flies carrying selection marker of 3xP3-RFP were further validated by genomic PCR and sequencing. CRISPR generates a 4,884-bp deletion removing the entire *NimA* CDS that is replaced by the cassette *Gal4-3xP3-RFP*.

**Table 1.** Fly list:

|  |  |  |
| --- | --- | --- |
| Fly |  |  |
| *srp(hemo)-3xmCherry* | Gift from D. Siekhaus<br>(Gyoergy et al., 2018) | Express RFP in macrophages. |
| *repo-nRFP 43.1* | (Laneve et al., 2013) | Express nuclear RFP in glia |
| *NimA-T2A-Gal4* | Build in this study. | *NimA KO-Gal4* allele |
| *UAS-mCD8RFP* | BDRC #32218 | Express membrane tethered RFP under UAS promoter. |
| *UAS-eGFP* | BDRC #5430 | Express enhanced GFP under UAS promoter. |
| *NrxGFP* | Gift from T. Volk<br>(Edenfeld, Volohonsky et al., 2006) | Neurexin IV – GFP fusion gene, marker of septate junction. |

### Bulk and single cell RNAseq analysis

The macrophage, glia and neuron bulk RNAseq data produced from E16 embryos and L3 larvae (wandering 3^rd^ larval instar) were uploaded from EBI database (accession: E-MTAB-8702, (Cattenoz, Sakr et al., 2020) and E-MTAB-14413 (Sakr, Cattenoz et al., 2022)). The raw data were mapped to the *Drosophila* genome August 2014 Dm6 (BDGF release 6 + ISO1 MT/Dm6) using Tophat (Trapnell, Pachter et al., 2009) and quantified with HT-seq (Anders, Pyl et al., 2015). For each stage, each tissue was compared to the two others, Log2 fold change and adjusted p values were estimated with DESeq2 (Anders & Huber, 2010). Raw read count and Log2 fold change are reported in **Supplementary Table S1**. Only genes differentially expressed significantly at both stages are represented on (**Figure 1A, Supplementary Figure S1A,B)**. GO term enrichment analysis (**Figure 1B**) was done using PANGEA (Hu, Comjean et al., 2023) using the whole gene set as background, the enriched terms are reported in **Supplementary Table S2**. The heatmaps (**Figure 1C, Supplementary Figure S1C**) were plotted in R (version 3.4.0) (R CoreTeam) using pheatmap (version 0.2) package (https://cran.r-project.org/web/packages/pheatmap/index.html).

The scRNAseq on adult CNS (**Figure 2A-A’’’**) were retrieved from (Davie et al., 2018) and the UMAPs were generated with the Seurat package using the FeaturePlot function (Hao, Hao et al., 2021, Stuart, Butler et al., 2019).

### Quantitative PCR

For the comparison between the expression levels of markers in macrophages and glia (**Supplementary Figure S1D**), cells were isolated from *srp(hemo)-3xmCherry* and *repo-nRFP* wandering L3, respectively. Staged lays of 3 h were carried out at 25°C and wandering larvae were collected 108–117 h after egg laying. L3 macrophages were isolated as mentioned in (Cattenoz et al., 2020). Larvae were bled in cold PBS containing PTU (Sigma-Aldrich P7629) to prevent macrophage melanization (Lerner and Fitzpatrick, 1950), and filtered (70 µm filter). For glia isolation, larval brains were dissected in cold PBS on ice and transferred to tubes containing 0.5 μg of collagenase IV (Gibco, Invitrogen) in 220 μL of PBS. Brains were incubated at 37°C on a Thermomixer with shaking at 500 rpm for 20 min. Then the brains were dissociated by pipetting up and down with 10-gauge needles and syringes and filtered through a 70 µm filter. Cells were sorted using FACS Aria II (BD Biosciences) at 4°C in three independent biological replicates for each genotype. Live cells were first selected based on the forward scatter and side scatter and only single cells were sorted according to the RFP signal. *Oregon-R* cells were used as a negative control to set the gate and sort the RFP positive cells, which were collected directly in TRI reagent (MRC) for RNA extraction. Around 100 000 cells were sorted for each replicate. The purity of the sorted populations was assessed by carrying out a post-sort step. The FACS sorter was set up to produce cell pools displaying at least 80% of purity on the post-sort analysis. The extracted RNA was treated with DNase I recombinant RNase free (Roche) and the reverse transcription was done using the Super-Script IV (Invitrogen) with random primers. The cycle program used for the reverse transcription is 65°C for 10 min, 55°C for 20 min, 80°C for 10 min. The qPCR was done using SYBR Green I Master (Roche). Actin5C (Act5C) and Ribosomal protein 49 (Rp49) were used to normalize the data. The primers are listed in the table below (**Table 2**). The p-values and statistical test used are indicated in the figure legends.

**Table 2:**
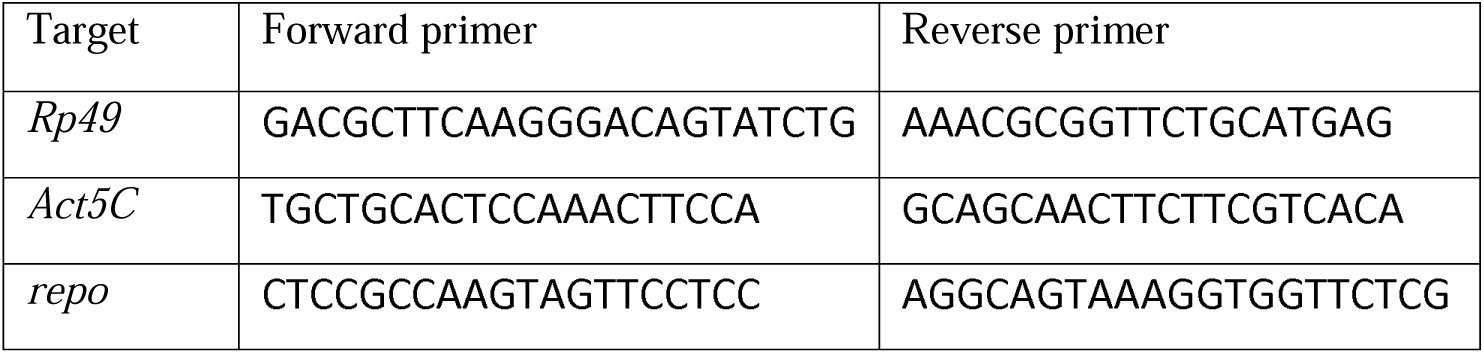
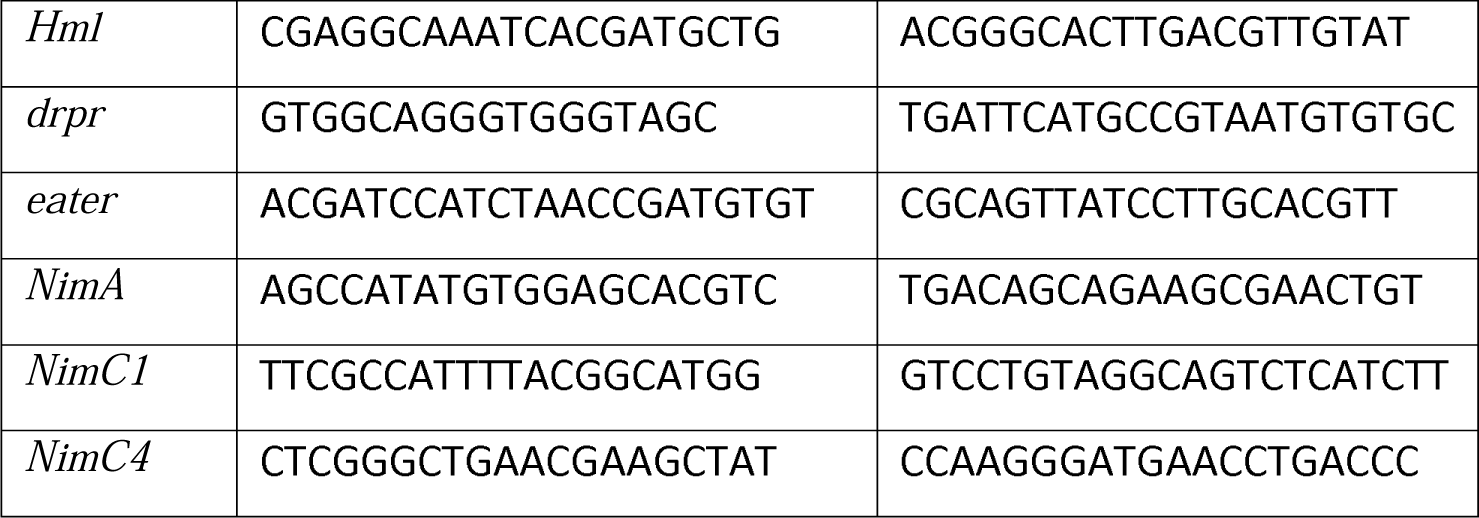
primer list.

### Immunolabeling and image analysis of CNS (embryo, larva, adult) and of *Drosophila* S2 cells

For L3 larvae and adults, CNSs from the indicated genotypes were dissected in cold PBS 1x and transferred into wells containing 4% PFA in 1xPBS and fixed for 2 h at RT. They were then washed in PTX (0.3% Triton x100 in 1x PBS) for 1 h and incubated in blocking reagent for 1 h at RT (O/N incubation at 4°C in blocking reagent for labeling with Dcp1 and deadpan). CNSs were then incubated with primary antibodies (**Table 3**) O/N at 4°C, washed in PTX 3x 10 min and incubated with secondary antibodies (Jackson) for 1 h at RT. After washing in PTX and labelled with DAPI, CNSs were mounted in Vectashield.

**Table 3:** list of antibodies.

| Antibody/labeling reagent | Ref | Dilution |
| --- | --- | --- |
| chicken anti-GFP | Abcam #ab13970 | 1/500 |
| rabbit anti-Nazgul | Gift from B. Altenhein | 1/100 |
| mouse anti-Repo | DSHB #8D12 | 1/20 |
| rat anti-RFP | Chromotek #5F8-150 | 1/500 |
| mouse anti-PDF | DSHB #C7 | 1/200 |
| mouse anti-cMyc | IGBMC antibody facility #Myc 9E10 | 1/200 |
| rabbit anti-HRP | Cappel #55974 | 1/500 |
| rat anti-Dpn | Abcam #ab195173 | 1/500 |
| mouse anti-pH3 | Millipore #05-806 (3H10) | 1/200 |
| rabbit anti-Dcp1 | Cell Signaling #9578S | 1/200 |
| rabbit anti-HA | Abcam #ab9110 | 1/500 |
| phalloidin Rhodamine | Cytoskeleton Inc. # PHDR1 | 1/200 |

Transfected *Drosophila* S2 cells were fixed 10 min with 4% PFA in 1x PBS at RT, washed 10 min in PTX at RT, incubated 1 h in blocking reagent at RT and incubated with the indicated primary antibodies diluted in blocking reagent O/N at 4°C. The following day, samples were washed 10 min in PTX at RT, incubated in secondary antibodies diluted 1:400 in blocking reagent for 1 h at RT and incubated in DAPI with or without phalloidin Rhodamine for 30 min at RT. Samples were finally washed in 1x PBS, mounted in Vectashield.

Images were acquired with Leica SP8 inverted-based confocal microscope or Leica spinning disk microscope. Image analysis was carried out using Fiji (National Institute of Mental Health, Bethesda, Maryland, USA, (Schindelin et al., 2012)). CNS volume as well as number of PH3/Dpn/Dcp1 positive cells in the CNS were estimated with the IMARIS software (RRID:SCR_007370) based on confocal images.

### Developmental delay/lethality and lifespan assay

To score for developmental delay and lethality, animals were let to lay eggs for 4 h at 25°C on apple juice agar plates supplemented with dry yeast. *w;NimA-T2A-Gal4* (*NimA KO-Gal4*, experimental genotype) or *w;NimA-T2A-Gal4/+* (control genotype, F1 from *NimA KO-Gal4* crossed with *w1118*) L1 larvae were collected 24 h after egg lay (AEL). For each genotype, 100 L1 were transferred in a fresh medium vial and let develop at 25°C. The number of newly formed pupae and eclosed adults was monitored daily. At least 9 vials (≥ 900 larvae) were analyzed per genotype.

To assess the lifespan, 1-day-old animals (*w;NimA-T2A-Gal4* or *w;NimA-T2A-Gal4/+)* were collected and transferred in a fresh medium vial so that each vial contained 10 males and 10 females. Vials were flipped 3 times per week for 60 days and the presence of dead flies was monitored at each flipping. 5 vials of 20 adults were analyzed per genotype (n=100).

### NimA expression vectors

To generate the vector *UAS-attb-NimA-T2A-mCD8GFP*, the NimA coding sequence was retrieved from *pOT2 NimA-RD* (DGRC #MIP14095) using primers containing NotI restriction site for the Forward primer targeting the 5’ end of NimA and AscI restriction site on the reverse primer targeting the codon stop of NimA coding sequence. The PCR was carried out using High fidelity Taq polymerase (Thermo Fisher Scientific). The mCD8GFP coding sequence was retrieved from genomic DNA extracted from the fly stock *QUAS-mCD8GFP* (BDRC #30001) using primers including AscI restriction site and the sequence coding for the self-cleaving peptide T2A on the forward primer and the restriction site KpnI on the reverse primer. The two PCR products were digested with their respective restriction enzymes and inserted in the *UAS-attb* vector (DGRC Stock 1419; https://dgrc.bio.indiana.edu//stock/1419; RRID:DGRC_1419) digested with NotI and KpnI.

For the vector *UAS-attb-NimA-Myc-T2A-mCD8GFP*, the Myc sequence was inserted in *UAS-attb-NimA-T2A-mCD8GFP* by PCR. To generate *UAS-attb-NimA-HA-T2A-mCD8GFP*, the Myc tag was replaced by the HA by PCR.

The *UAS-attb-T2A-mCD8GFP* vector was produced by inserting the *T2A-mCD8GFP* sequence in *UAS-attb* empty vector.

The sequence of each plasmid was verified by sanger sequencing.

### Phagocytosis and aggregation assays

For phagocytosis assay, six million *Drosophila* S2 cells were plated per well in a 6-well plate with 1.5 mL of Schneider medium +10% FCS+0.5% PS. Transfections were carried out 12 h after plating, using the Effectene Transfection Reagent (Qiagen) (Cattenoz, Popkova et al., 2016). 1 µg of Gal4 expression vector (SKactin-Gal4, RRID:DGRC_1019) was co-transfected with 1 µg of vector *UAS-NimA-T2A-mCD8GFP, UAS-NimA-Myc-T2A-mCD8GFP* or *UAS-T2A-mCD8GFP* (control). 48 h after transfection, the cells were collected and plated in 96 well plate (100 µL per well) and challenged with RFP-fluorescent latex beads (polysciences #19507-5) diluted 1/500 for the indicated time (0 or 20 min). The cells were then rinsed twice with PBS, fixed with 4% PFA in 1x PBS for 10 min. Images were acquired for each condition by confocal microscopy and the RFP intensity was estimated using Fiji.

To estimate the aggregation of the cells, three million *Drosophila* S2 cells were plated per well in a 6-well plate with 1.5 mL of Schneider medium +10% FCS+0.5% PS. Transfections were carried out 1 h after plating, using the Effectene Transfection Reagent (Qiagen #301427) according to the manufacturer’s instructions. 0.5 µg of Gal4 expression vector (SKactin-Gal4, RRID:DGRC_1019) was co-transfected with 0.5 µg of vector *UAS-attb-NimA-HA-T2A-mCD8GFP* (*NimA-HA GOF*) or 0.5 µg of vector *UAS-attb-T2A-mCD8GFP* (Control). Transfection was carried out for 48 h at 28°C. Cells were then fixed with 4% PFA in 1x PBS, immunolabelled with anti-GFP and DAPI as described above and scanned using CellInsight CX7 instrument (Cellomics) at 20x magnification (60 fields per well). The GFP signal was used to determine the boundary of the clumps and DAPI was used to count the number of nuclei per clumps. The signal was estimated using HCS Studio software and Colocalisation Bioapplication.

### Blood-Brain Barrier permeability assay

Larvae *w1118* and *NimA-T2A-Gal4/+* were used as controls and compared to homozygous *NimA-T2A-Gal4* (*NimA KO-Gal4)*. Animals were staged with 4 h egg lay at 25°C. L1 were collected 24 hrs after egg laying (AEL). For each genotype, 100 L1 were transferred in a fresh medium vial and incubated at 25°C until wandering L3 stage. L3 were dissected in cold PBS 1X and incubated for 30 min in 0.25 mM of 10 kDa dextran Texas red (Invitrogen). The CNSs were washed 2 times of 10 min in 4% PFA at room temperature (RT) with agitation (50 rpm) to remove excess dextran, followed by incubation in 4% PFA for 1 hour and 40 minutes. The CNS were mounted in Vectashield and acquired using Leica spinning disk microscope with 63x objective and 1μm confocal section. The dextran intensity was measured using Fiji (National Institute of Mental Health, Bethesda, Maryland, USA, (Schindelin et al., 2012).

### *In situ* hybridization

We used a RNAscope® probe targeting NimA (RNAscope Dm-NimA-C3 #1082391-C3) and reagent kit. Larval CNSs from *Oregon R* were dissected in cold PBS 1x and fixed in 4% PFA O/N at 4°C then washed 3x 10 min in PBST (0.1% Tween-20 in 1x PBS) followed by one wash in PBS 1x. Samples were then incubated in RNAscope reagents as in protease III at 40°C for 10 min and washed in 3x 2 min in PBS 1x. The probe targeting *NimA* was then added in addition to the positive control probe and samples were incubated O/N at 40°C. The amplifiers and detection solution in the RNAscope reagent kit were added sequentially for hybridization signal amplification for the indicated time. CNSs were then mounted in Vectashield, imaged using Leica spinning disk microscope and analyzed using Fiji.

### Co-immunoprecipitation (IP) and immunoblot analysis

3.10^6^ *Drosophila* S2 cells were plated per well in a 6-well plate with 1 mL of Schneider medium +10% FCS+0.5% PS. Transfections were carried out 1 h after plating, using the Effectene Transfection Reagent (Qiagen #301427) according to the manufacturer’s instructions. For the “NimA-HA, NimA-Myc GOF” sample, cells were co-transfected with *SKactin-Gal4, UAS-attb-NimA-HA-T2A-mCD8GFP* and *UAS-attb-NimA-Myc-T2A-mCD8GFP*, 0.3 µg each. Control cells were co-transfected with *SKactin-Gal4* and *UAS-attb-T2A-mCD8GFP*, 0.5 µg each. Transfections were carried out for 72 h at 28°C.

At least 6.10^6^ transfected cells were used for Co-IP experiments. Cells were washed in 1x PBS and resuspended in 500 µL of lysis buffer (50umM Tris–HCl pH 7.5, 150 mM NaCl, 10% glycerol, 1 mM EDTA, 1% Triton, 1x proteinase inhibitor cocktail (Roche). Samples were incubated 10 min on ice, then centrifuged at 10000 x g for 5 min at 4°C. 10% of the cell lysate was used as Input, complemented with 2X SDS buffer containing 0.2 M DTT and boiled at 95°C for 5 min. The rest of the cell extract was used for IP with anti-HA agarose beads (Monoclonal anti-HA-agarose antibody produced in mouse Sigma #A2095). Before use, beads were washed three times in lysis buffer and resuspended in 500 µL of lysis buffer. A “bead control” sample was included in each experiment upon incubating the beads with 950 µL of lysis buffer. IP samples were incubated O/N at 4°C in agitation, washed three times in lysis buffer, eluted in DTT-free, 2X SDS buffer and boiled at 95°C for 5 min. Beads were removed by centrifugation and 0.1 M DTT was added. INPUT and IP samples were then processed by western blot.

Proteins were separated with 10% SDS–PAGE gel, transferred onto nitrocellulose membrane and probed with primary antibodies diluted in 2% milk in 1x PBS. The following antibody were used: mouse anti-Myc (1:1000), mouse anti-HA (1:1000), mouse anti-Tubulin (1:2000), chicken anti-GFP (1:1000). Signals were detected using anti-mouse and anti-chicken HRP-conjugated antibodies (1:5000) and the chemiluminescent substrate SuperSignal West Pico PLUS (Thermo Fisher Scientific *#*34580). Chemiluminescence detection was performed Amersham Imager 600.

### Septate junction analysis with *NrxGFP* animals

Larvae *NrxIVGFP* and *NimA-T2A-Gal4/+;NrxIVGFP* were used as controls and *NimA-T2A-Gal4;NrxIVGFP* as *NimA KO-Gal4*. Animals were stage with 4 h egg lay at 25°C. L1 were collected 24 hrs after egg laying (AEL). For each genotype, 100 L1 were transferred in a fresh medium vial and incubated at 25°C until wandering L3 stage. The L3 were dissected in cold PBS 1X carefully without touching the CNSs. Then, CNSs were incubated for 30 minutes in 4% PFA at room temperature (RT) on the shaker. The CNSs were washed three times for 10 min with 0.3% PTX, incubated with blocking reagent for 1 hr at RT on the shaker, incubated overnight at 4°C with primary antibodies, washed three times for 10 min with 0.3% PTX, incubated for 1 hr with secondary anti-bodies, washed two times for 10 min with PTX, incubated for 40 min with DAPI and then mounted in Vectashield. The following of primary antibody was used to label NrxGFP: Chicken anti-GFP (1/1000). Secondary antibody used was: donkey anti-chicken coupled with FITC (1/400). The central area of brain lobes was imaged using Leica spinning disk microscope with 63x oil objective with 0.2 μm confocal section. The junctions were analysed using Imaris (Version 10.1.1) to reconstruct the BBB junctions. The filament statistics, number of segments, and filament volume were measured on the 3D reconstruction for NrxGFP at the surface of the CNS optic lobes. Then, the ratios of [(number of segments -1) / filament length] (*i.e.* gap per µm) and [volume of the filament / filament length] (*i.e.* average cross-section area) were determined for each lobe.

## Acknowledgments

We thank Lara El Berjawi, Valentine Jungmichel, Luca Sartori, Alexia Pavlidaki, Maria Dolores De Donno for technical help.

We acknowledge the Imaging Center of the IGBMC, member of the national infrastructure France-BioImaging supported by the French National Research Agency (ANR-10-INBS-04). The sequencing was performed by the GenomEast platform, a member of the “France Génomique” consortium (ANR-10-INBS-0009). We thank A. Maglott-Roth from the IGBMC screening facility for the macrophage analysis. We thank D. Siekhaus and B. Altenhein for providing fly stocks and antibodies. In addition, stocks obtained from the Bloomington *Drosophila* Stock Center (NIH P40OD018537) and antibodies obtained from the Developmental Studies Hybridoma Bank created by the NICHD of the NIH and maintained at The University of Iowa (Department of Biology, Iowa City, IA 52242) were used in this study. This work was supported by Inserm, CNRS, UDS, Ligue Régionale contre le Cancer, Hôpital de Strasbourg, ARC, CEFIPRA, USIAS, FRM and ANR grants, and by the CNRS/University/Inserm IRP Machub. R. Sakr was supported by the French state fund through a doctoral contract from the University of Strasbourg, the Fondation pour la Recherche Médicale (FDT2020010107630) and the ANR. S. Monticelli was funded from CEFIPRA, FRC and ANR. G. Zhang was funded from the Chinese Scholarship Council. T. Tabiat was funded by the ANR. This work of the Interdisciplinary Thematic Institute IMCBio, as part of the ITI 2021-2028 program of the University of Strasbourg, CNRS and Inserm, was supported by IdEx Unistra (ANR-10-IDEX-0002), and by SFRI-STRAT’US project (ANR 20-SFRI-0012) and EUR IMCBio (ANR-17-EURE-0023) under the framework of the French Investments for the Future Program.

## Figure legends

**Supplementary Figure S1: Comparison of bulk RNAseq and scRNAseq of E16 and L3 macrophages**

**A,B**) Comparison of the transcriptomes of neurons, macrophages and glia in stage 16 embryos (E16) and wandering larvae (L3). The scatter plots represent the Log2 fold change (L2FC) of differentially expressed genes enriched in neurons (**A**) and macrophages (**B**) compared to the two alternative tissues (i.e. neurons vs. macrophages and glia in **A**, macrophages vs. glia and neurons in **B**). Only genes presenting significant fold change in both E16 and L3 were plotted, the x-axes represent the L2FC in E16 and the y-axes the L2FC in L3. The complete data are available in **Supplementary Table S1**.

**C)** Heatmap representing the expression levels of known and new markers of neurons, macrophages and glia. The color gradient is representative of the expression levels from blue to red, low to high expression.

**D)** Expression levels of the indicated genes in glia and macrophages estimated by qPCR, n=4, p-value estimated by bilateral student test for equal variance, y-axis in Log10 scale.

**Supplementary Figure S2:Expression profile of *NimA-T2A-Gal4* in the adult CNS**

**A)** Annotation of the ellipsoid body and the mushroom body lobe in the adult CNS. The image is from the Virtual Fly Brain Atlas (Bogovic, Otsuna et al., 2020, Court, Costa et al., 2023).

**B,C)** Immunolabelling of adult CNS *NimA-T2A-Gal4/+;UAS-eGFP/+* with anti-Repo (in white), anti-Naz (in red) and anti-GFP (in green), nuclei were labelled with DAPI. Single sections are shown and highlight the expression of NimA in Naz positive glia (ALG) as well as Naz negative glia (EG) surrounding the ellipsoid body and the mushroom body. The scale bars represent 20 µm.

**Supplementary Figure S3: Expression profile of *NimA-T2A-Gal4* in the larval CNS and PNS**

**A)** *NimA* expression profile in larval brain using fluorescent *in situ* hybridization. The whole CNS is shown in (**A**) and higher magnifications focus on the optic lobe (**A’**) and the ventral nerve cord neuropil (**A’’)**, a cross-section is shown on the right panel. *NimA* is shown in green and DAPI was used to label the nuclei (in blue). The scale bars represent 100 µm or 20 µm as indicated on the images.

**B-D)** Immunolabelling of ventral nerve cord (**B,C**) and peripheral nerves (**D**) from *NimA-T2A-Gal4/+;UAS-mCD8RFP/+* L3 with anti-HRP (in green), anti-Repo (in white) and anti-RFP (in red). The tract ensheathing glia (TEG), an ensheathing glia subtype that wrap the nerves exiting the VNC, are shown by a white arrowhead and the wrapping glia (WG) are indicated by white lines. Scale bars are 100 µm, 50 µm and 25 µm.

**E-F)** Immunolabelling of L3 CNS *NimA-T2A-Gal4/+;UAS-eGFP/+* with anti-Naz (in red), anti-Repo (in white) and anti-GFP (in green), nuclei were labelled with DAPI. Subperineural glia (SPG) and perineural glia (PG) are highlighted in the BBB around the optic lobe (**E-E’’**), note that only the SPG express NimA. Ensheathing glia (EG) and astrocyte-like glia (ALG) are shown in the ventral nerve cord (VNC) (**F-F’’’**). Scale bars represent 50 µm, the left panels show the overlay of the four channels, single channels are shown in the successive panels.

**Supplementary Table S1: raw read counts and differential expression comparison of bulk RNAseq from E16 and L3 macrophages, glia and neurons.**

The first sheet of the table contains the raw read counts, the second sheet contains the differential expression values (L2FC) generated with DESeq2 comparing for each stage, macrophages versus glia and neurons (HvsNG), glia versus macrophages and neurons (GvsNH) and neurons versus glia and macrophages (NvsGH). The Log2 fold change (L2FC) and the adjusted p-value are reported for each gene and the FBGN ID as well as the gene symbol are indicated. The last column of the table highlights the markers of each tissue.

**Supplementary Table S2: GO term enrichment analysis carried out with PANGEA on the markers of each tissue.**

