## Supplementary figures and images for "NimA promotes cell adhesion at the blood brain barrier of *Drosophila* nervous system"

### Supplementary Figures

#### Supplementary Figure S1

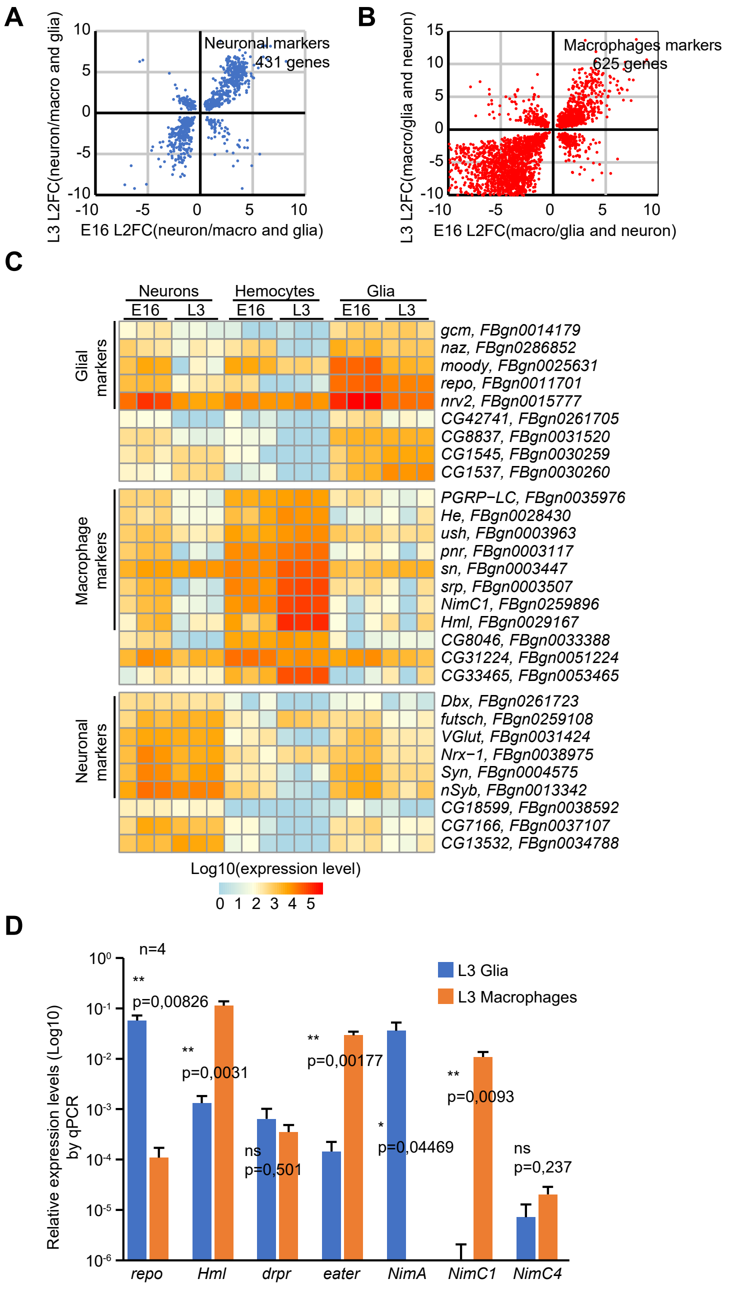

#### Supplementary Figure S2

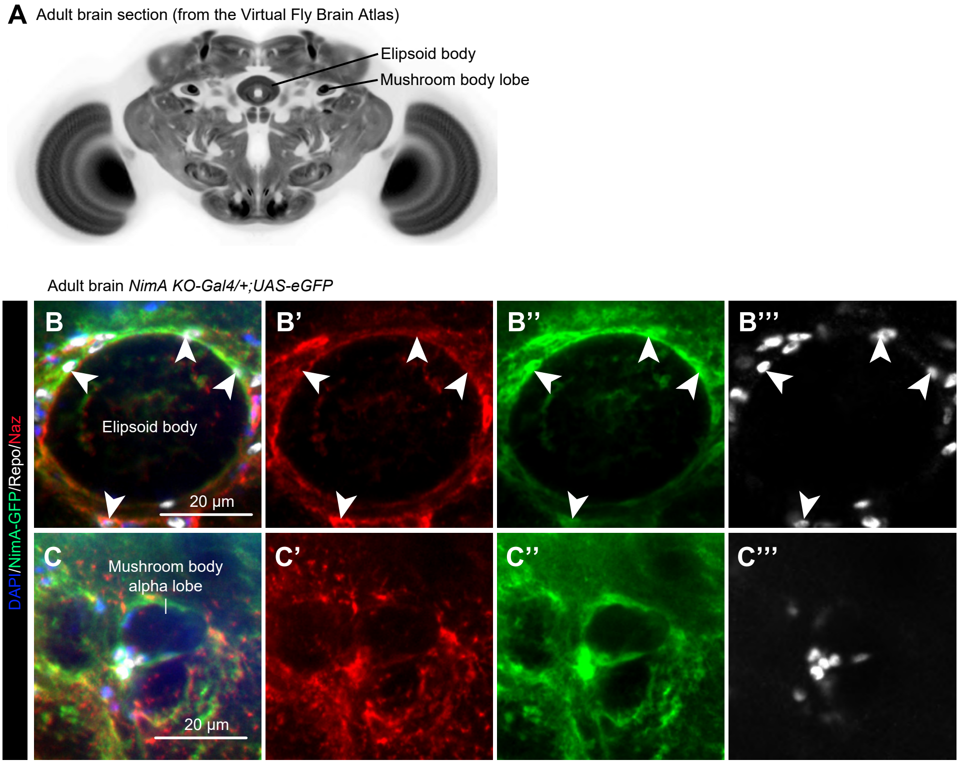

#### Supplementary Figure S3

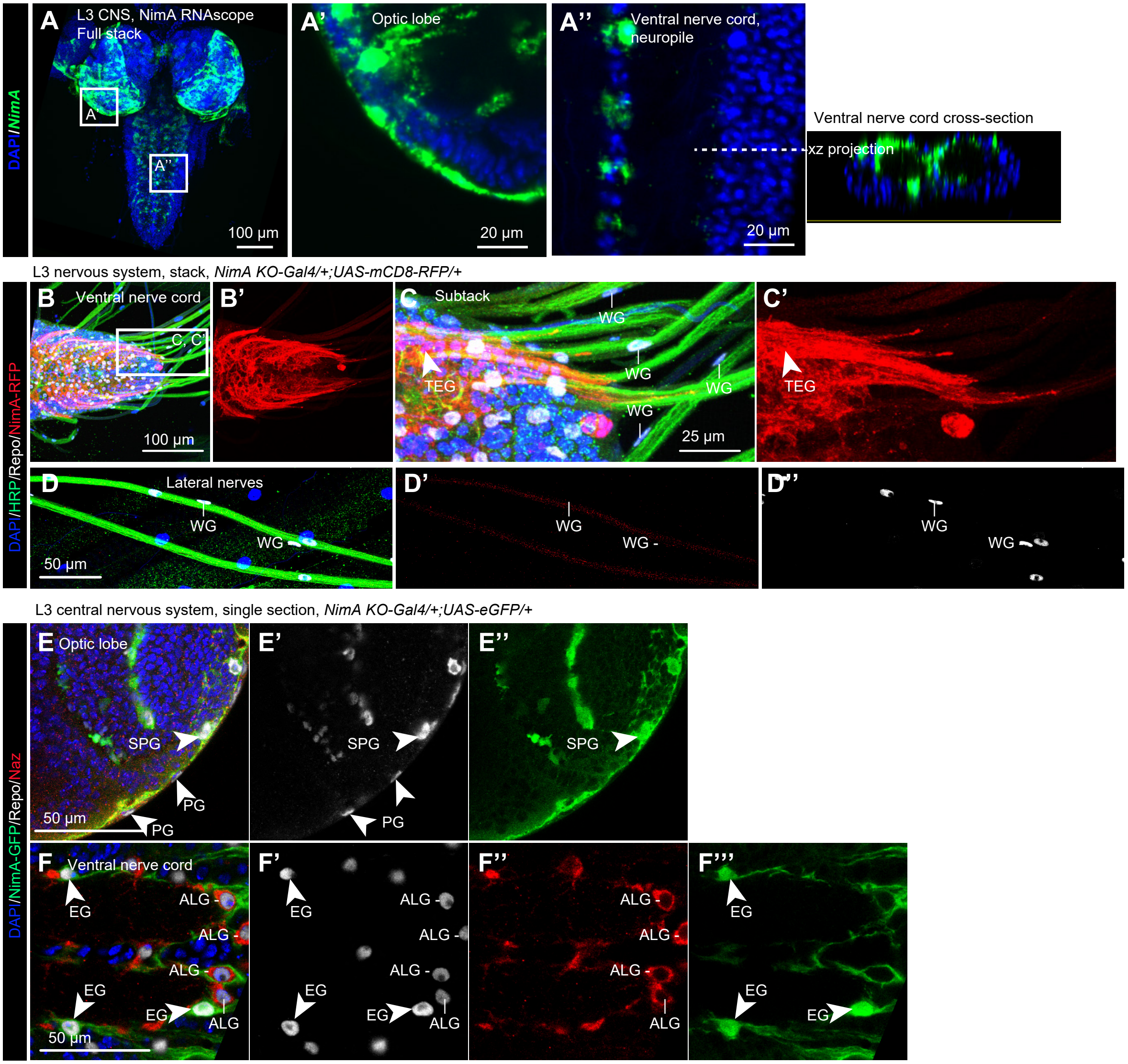
